# A Bayesian Nonparametric Approach to Discover Clinico-Genetic Associations across Cancer Types

**DOI:** 10.1101/623215

**Authors:** Melanie F. Pradier, Stephanie L. Hyland, Stefan G. Stark, Kjong Lehmann, Julia E. Vogt, Fernando Perez-Cruz, Gunnar Rätsch

**Affiliations:** Computational Biology Program, Memorial Sloan Kettering Cancer Center, New York, U.S.A.; University Carlos III in Madrid, Leganés, Spain; School of Engineering and Applied Sciences, Harvard University, Cambridge, USA; Ph.D. Program in Computational Biology and Medicine, Weill Cornell Medicine, New York, U.S.A.; Department of Computer Science, ETH Zürich, Zürich, Switzerland; Medical Informatics Group, University Hospital Zürich, Zürich, Switzerland; Department of Mathematics and Computer Science, University of Basel, Basel, Switzerland; Swiss Institute for Bioinformatics, Lausanne, Switzerland; Swiss Data Science Center, ETH Zürich and EPFL Lausanne, Switzerland; Department of Biology, ETH Zürich, Zürich, Switzerland

## Abstract

**Motivation:** Personalized medicine aims at combining genetic, clinical, and environmental data to improve medical diagnosis and disease treatment, tailored to each patient. This paper presents a Bayesian nonparametric (BNP) approach to identify genetic associations with clinical/environmental features in cancer. We propose an unsupervised approach to generate data-driven hypotheses and bring potentially novel insights about cancer biology. Our model combines somatic mutation information at gene-level with features extracted from the Electronic Health Record. We propose a hierarchical approach, the hierarchical Poisson factor analysis (H-PFA) model, to share information across patients having different types of cancer. To discover statistically significant associations, we combine Bayesian modeling with bootstrapping techniques and correct for multiple hypothesis testing.

**Results:** Using our approach, we empirically demonstrate that we can recover well-known associations in cancer literature. We compare the results of H-PFA with two other classical methods in the field: case-control (CC) setups, and linear mixed models (LMMs).

## 1 Introduction

Cancer encompasses not one, but a large group of genetic diseases involving abnormal cell growth with the potential to invade or spread to other parts of the body. Although a small set of universal underlying principles were identified, the so-called “hallmarks” of cancer [19, 20], each type of cancer presents unique properties, making this disease very hard to treat [37, 22, 43].

Genetic association studies have been successful in relating somatic mutation to carcinogenesis, but has been limited to the detection of common large-effect variants in the presence of only small cohorts [9, 27, 8]. Finding somatic driver mutations is even more challenging since these mutations are often rare. The large phenotypic heterogeneity, which reduces statistical power in the discovery method, causes some associations to remain hidden [38, 29, 40]. Cohort sizes tend to be small, especially in rare cancers, which makes the discovery of small effect size associations difficult [3]. Additionally, carcinogenesis is driven by the accumulation of mutations that may act epistatically or pleiotropically during the disease, further reducing the power of typical approaches [48, 11, 42, 12]. To overcome these difficulties, new approaches for interpreting genetic variation across different cancer types are required.

In recent years, efforts to mine electronic health records (EHRs) show promise to impact nearly every aspect of healthcare [26]. The adoption of EHRs in hospitals has increased dramatically and has become a powerful resource for phenotyping [2, 32], with the potential for establishing new patient-stratification principles and for revealing unknown disease correlations. Integrating EHRs data with genetic data will also give a better understanding of genotype-phenotype relationships [49]. EHRs consist of both structured and unstructured information. Structured data is a valuable source of information that includes billing codes, laboratory reports, physiological meamelements, and demographic information, among others. Yet, most of the clinical data comes as unstructured notes, e.g., around 98% of the EHRs [26]. These include a broad spectrum of clinically-relevant information which might be useful to identify novel phenotypic relationships so far unknown by the clinicians [32, 26].

This work presents a joint generative model to discover associations between somatic mutations and clinical features in cancer that deals with phenotype heterogeneity, small cohort size, epistasis and pleiotropy in a straightforward way. Our method infers latent topics from the clinical text and genetic information, capturing complex interactions between groups of genes and clinical features. It is directly inspired by the Poisson factorization model for recommendation systems [16], with three important differences.

First, we introduce confounding effects as conditional variables, i.e., variables that might cause spurious associations to appear. In particular, our model considers multiple types of cancer together (the type of cancer is treated as a confounder) and shares information among all patients in a hierarchical fashion. Indeed, most cancers are known to share common pathogenesis despite specificities of the cell type and tissue origin [43]. By doing so, specific effects for each type of cancer can be isolated, and additional (less well-known) associations with somatic mutations of smaller effect size can be obtained.

Second, we force sparsity on the textual and genetic topics by using shape parameters smaller than one in the Gamma distribution priors. Sparsity is crucial to find meaningful, easy-to-interpret associations; those can be validated either through previous studies by looking in the literature, or subsequent tests in the lab.

Third, we present a nonparametric alternative model to [16] by replacing the continuous patient weights with a binary matrix whose probability distribution is induced by a hierarchical extension of the Indian buffet process (IBP). Also in the literature, the authors in [17] propose a nonparametric Poisson factorization model, but they rely on a stick-breaking construction different from the IBP, and the weights are continuous, which renders interpretability of the latent variables more tedious. The discrete nature of the IBP helps in terms of interpretability and allows combining the proposed Bayesian model with classical frequentist approaches for statistical testing between the inferred patient partitions. An efficient Markov chain Monte Carlo (MCMC) procedure based on a slice sampler for the hierarchical IBP is presented.

Bayesian modeling has already been proven useful for epistasis [53], pleiotropy [53, 52] or sub-phenotyping applications [35, 28]. To our knowledge, the proposed model is the first one to deal with clinical text data and genetic information jointly in a Bayesian nonparametric way, capturing phenotypic heterogeneity, epistasis and pleiotropy in a straightforward way while correcting for the cancer type as confounder. We consider multiple cancers jointly in order to increase statistical power, allow for the analysis of rare cancers, and identify fundamental mechanisms shared across different types of cancer.

## 2 Methods

### 2.1 Study design and setting

The study was designed as a retrospective cohort study for the development and analysis of techniques to analyze clinical narratives in the context of somatic mutations. It was performed at Memorial Sloan Kettering Cancer Center (MSKCC). The institutional review board of MSKCC provided a Waiver of Authorization (WOA; WA0426-13) for this study. Clinical notes were provided by the IT services group at MSKCC. The Center for Molecular Oncology at MSKCC provided the information about somatic mutations from the MSK Impact panels of patient tumors. We included 1,946 patients for which we had MSK Impact panel data and at least one clinical narrative available at the time of delivery. Data analyses were performed on the HPC compute systems at MSKCC. Additional statistical analyses were performed at University Carlos III and ETH Zürich.

### 2.2 Database Description

So far, genomic testing of tumors has been done routinely only for certain solid cancer tumors, such as melanoma, lung, or colon cancer. For most cancers, the available tests have been limited to analyzing one or a handful of genes at a time, and within each gene, only the most common mutations could be detected.^1^ A new targeted tumor sequencing test called MSK-IMPACT (Integrated Mutation Profiling of Actionable Cancer Targets) is able to detect somatic mutations and other critical somatic aberrations in both rare and common cancers [10, 51].

Using the MSK-IMPACT panels[51], somatic mutations regarding specific screened genes can be obtained as follows. For each patient, tumor cells are compared with healthy cells of that same patient, extracted from the blood stream, as illustrated in Figure 1. In this work, a gene is said to be mutated when there exists at least one difference in the sequence between the tumor cells and healthy cells for that particular gene.^2^ We finally obtain a binary matrix for *N* = 1946 patients and *G* = 410 genes where “1” encodes for a mutated gene and “0” otherwise.

**Figure 1:**
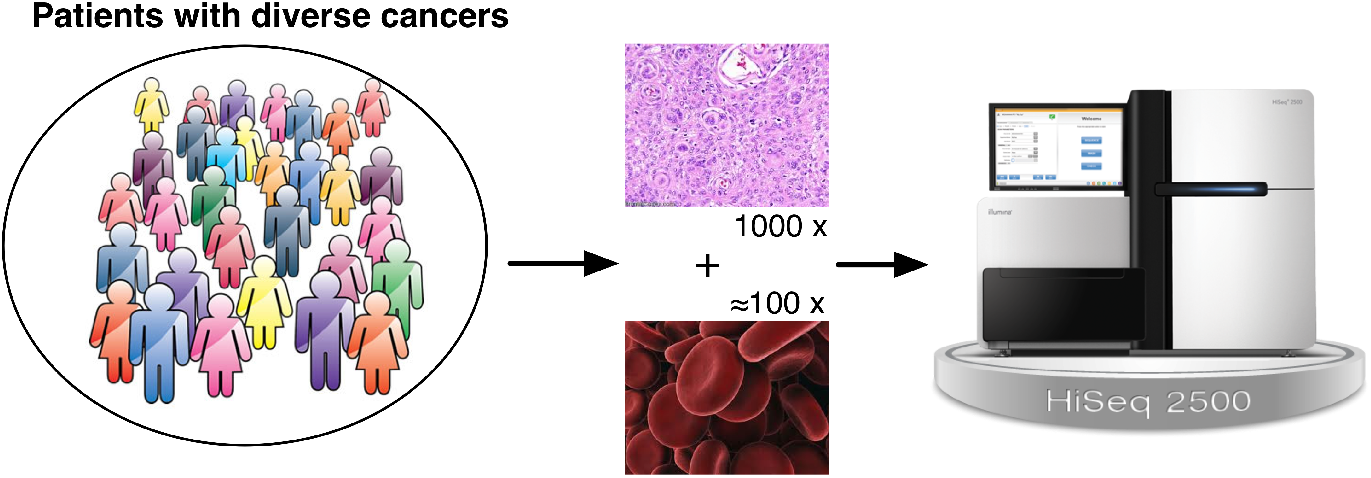
Diagram illustrating molecular profiling of tumors with the MSK-IMPACT Panel. (top) For each patient, tumor DNA is compared against normal tissue DNA in order to detect somatic mutations. The MSK-IMPACT panel is routinely used for a large number of patients per year at MSKCC [51].

The screened genes have been shown to play a role in the development or behavior of tumors, although their individual relation to specific phenotypes remains obscure [10, 51].

Concerning the clinical information, based on all EHRs, we build a bag-of-word representation of unified medical language system (UMLS) terms, extracted using the *Metamap*^3^ processing tool [4]. The UMLS refers to a standardized, comprehensive thesaurus and ontology for biomedical concepts, whose objective is to provide facilities for natural signal processing tasks [7]. Since each patient can have a varying number of records, we group all clinical history into a single EHR, and only consider the appearance or absence of each UMLS term, i.e., binarized clinical features. We compute the tf-idf score for each UMLS term, and only keep the 300 clinical terms with highest score.

The database includes clinical and genetic information for *N* = 1946 patients and 5 different cancer types: bladder cancer, breast carcinoma, colorectal cancer, non-small cell lung cancer, and prostate cancer. We consider genes and UMLS terms that are present in at least 1% of the patient population, resulting in *D* = 249 dimensions, including 72 genes and 177 clinical terms.

Even if the dataset is binary, we can use a Poisson likelihood because of the high sparsity degree of such matrix (7.28% of non-zero values). In such scenario, the Poisson distribution is a good approximation of a Bernoulli, and we adopt it by mathematical convenience in the inference process [14, 46]. In the following, we use this data for: a) discovering latent factors and, b) testing for significant features (either genetic or phenotypical) associated to each factor.

### 2.3 Classical Approach: Case-Control Setup

The most common approach for genetic association studies is the case-control (CC) setup, which compares two large groups of individuals, one case group presenting a particular phenotype, and one control group without such phenotype.All individuals in each group are genotyped to identify somatic mutations in a panel of genes. For each of these genes, it is then investigated if the somatic mutation is significantly associated with the phenotype of interest. In such setups, the fundamental unit for reporting effect sizes is the odds ratio. The odds ratio in this case refers to the odds of exhibiting the phenotype for individuals having a specific somatic mutation and the odds of exhibiting the phenotype for individuals who do not present such somatic mutation. A *p*-value for the significance of the odds ratio is typically computed using a simple *χ*-squared test or Fisher test. Finding odds ratios that are significantly different from one is the objective of an association study because this shows that there is statistical evidence that the somatic mutation is associated with the phenotype.

### 2.4 Confounder Correcting Approach: Linear Mixed Model

Linear mixed models (LMMs) have proved particularly useful for genetic association studies due to its capacity to account for confounding effects and limit the number of false associations [31, 30]. Let **X** be the observation matrix, where each element *x_ng_* corresponds to an indicator variable for a particular patient *n* ∈ {1,…, *N*} and somatic mutation in gene *g* ∈ {1,…, *G*}, and *x_g_* ∈ {0,1}^*N*×1^ is the indicator vector for gene *g* across all patients. The binary variable *x_ng_* indicates whether any somatic mutation occurred in the corresponding gene. Let *y_nq_* be the binary indicator variable of the presence of a certain clinical feature *q* ∈ {1,…, *Q*} for a given patient *n*, and ***y***_*q*_ ∈ ℝ^*N*×1^ the indicator vector for clinical feature *q* across all patients. Finally, let us define *c_nl_* as the binary indicator variable of patient *n* to the cancer type *ℓ* ∈ {1,…,*L*}, and ***c***_*n*_ ∈ {0,1}^1×*L*^ the cancer type assignment vector, where Σ*_ℓ_ c_nℓ_* = 1 (for simplicity, we only consider patients having one single type of cancer). For each pair of gene *g* and clinical feature *q*, a LMM can be defined as follows:

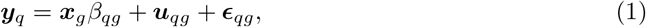

where *β_qg_* ∈ ℝ refer to the fixed effect of feature *q* and gene *g*, and ***u***_*qg*_, *ε_qg_* ∈ ℝ^*N*×1^ are the random effects (structured noise and observational noise, respectively). The prior assumptions for the structured and uniform noises are 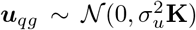 and 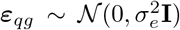, where **K** refers to a similarity matrix between the patients, for instance, the cosine similarity of the cancer type assignment vectors ***c***_*i*_ and ***c***_*j*_, **K** = **CC**^*T*^. The LMM assumes that the output ***y_q_*** is Gaussian-distributed:

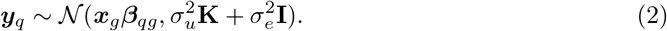

When the data is binary or count data, a common practice is to apply a standard rank-based inverse normal transformation beforehand as a preprocessing step [24], although this has become controversial more recently [5]. LMMs are discriminative since they try to model the conditional probability *p*(***y***_*q*_|***x***_*g*_). In this paper, we propose an alternative generative approach that models the joint distribution *p*(***y***_*q*_, ***x***_*g*_) and captures complex correlations via latent factors.

### 2.5 Hierarchical Bayesian Nonparametric Approach

Let 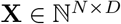 be the observation matrix of count data for *N* patients and *D* dimensions, where *D* includes both clinical and genetic information, i.e., *D* = *G* + *Q*, where *G* is the number of genes and *Q* is the number of clinical terms. In the following, we propose two Poisson factor analysis (PFA) approaches to model the joint observation matrix **X** of genetic information and clinical data. In these models, patients will be represented by binary feature activation vectors, and each of these features will capture common correlation patterns among the somatic mutations and clinical term occurrences.

#### 2.5.1 Poisson Factor Analysis (PFA)

We first consider a nonparametric non-negative matrix factorization model with Poisson likelihood and Gamma-distributed factors:

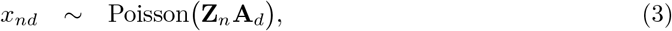

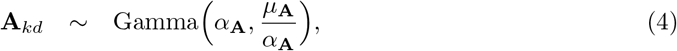

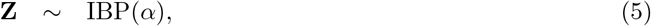

where *α* is the concentration parameter of the IBP controlling the a priori number of ones in matrix **Z** (i.e., the a priori expected number of latent features), and *μ*_**A**_, *α*_**A**_ are the prior mean and shape parameter for each element of matrix **A.** Sparsity can be induced easily in the factors by choosing *α*_**A**_ ≪ 1. Inference is performed using an MCMC approach based on a semi-ordered stick-breaking representation of the IBP prior [44].

#### 2.5.2 Hierarchical Poisson Factor Analysis (H-PFA)

Although different types of cancer are known to share similar phenotypes and underlying mechanisms (shared activation of certain pathways), the mutation rate and phenotype occurrence might vary in different proportions, according to each type of cancer. Given this premise, we propose a hierarchical Bernoulli process Poisson factor analysis model to allow for different feature activation levels depending on each type of cancer. In the following, we will shorten the name of this model to hierarchical Poisson factor analysis (H-PFA).

Let *r_n_* ∈ [1,…, *L*] be a categorical variable indicating the type of cancer of patient *n* among the total number of cancer types *L* (*r_n_* corresponds to the index of the non-zero value in vector ***c***_*n*_ defined in Section 2.4). A hierarchical construction can be formulated based on the finite representation of the IBP and letting *K* → ∞, such that different levels of feature activation are allowed for each type of cancer. Let *ρ_k_* be the global activation probability of feature *k*, and 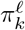 be the specific activation probability of feature *k* for cancer type *ℓ* ∈ [1,…, *L*]. We can then assume that each specific activation probability is Beta-distributed such that 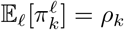:

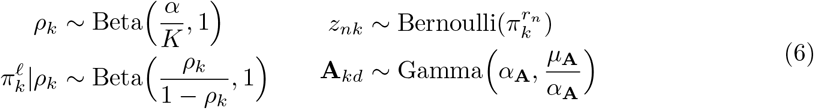

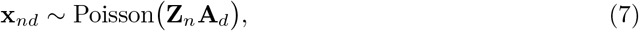

where the feature activation variables in vector **Z**_*n*_ are drawn from different activation probability vectors {***π***^1^,…, ***π***^*L*^} depending on the type of cancer *r_n_* of patient *n*. When *K* → ∞, this prior over **Z** is equivalent to a hierarchical Beta process (BP) construction [45] on top of Bernoulli processes (BePs) in the De Finetti representation. In the same way that a hierarchical Dirichlet process (HDP) allows for atom sharing with varying weights across different groups of data, the H-PFA allows for feature sharing with different activation weights across different types of cancer.

### 2.6 Statistical Methodology

Once the model has been trained (samples from an approximate posterior distribution can be drawn), we proceed with a classical frequentist approach^4^ to identify statistically significant clinico-genetic associations across cancer types. First, we take *M* posterior samples from the posterior distribution of **Z** given **A** fixed. For each sample, patients that have the same feature assignment vector (activation pattern of features) can be grouped together in the same subpopulation. For instance, subpopulation (1001) refers to all patients having the first and fourth feature active. Let *P* refer to the total number of inferred subpopulations across the *M* posterior samples. By considering multiple posterior samples, we obtain slightly different partitions of patients in subpopulations. This can be seen as performing *soft-clustering* of patients, i.e., patients that are in-between subgroups might be assigned to different subpopulations in different posterior samples. Thus, the method is more robust against model inaccuracies at clustering patients. This is an important benefit of the Bayesian framework.

Next, to make our method robust against outliers (e.g., patients with rare features), we perform bootstrapping *B* times for each subpopulation and posterior sample. Bootstrapping relies on random sampling with replacement. It is a technique used for computing robust estimators against outliers by sampling from an approximating distribution, which is particularly useful for hypothesis testing when the model assumptions are in doubt or unknown [50]. The standard bootstrapping approach relies on the construction of an estimator for hypothesis testing based on a number *B* of resamples with replacement of the observed dataset (and of equal size to the observed dataset), i.e., sampling with replacement from the empirical distribution of the observed data.

Finally, given *M* posterior samples and *B* bootstrapping instances for each sample, we end up with *MB* different subpopulation instances. Measures of effect size (quantitative measure of the difference between two subpopulations) and statistical significance can be computed for each instance and then averaged across them, so that partition inaccuracies and outlier effects are mitigated. To identify statistically significant dimensions for each latent feature *k* = 1,…,*K* in sample *m* and bootstrap *b*, we split the whole patient population in two subgroups, 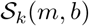 and 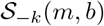, corresponding to patients whose latent feature *k* is active or inactive respectively, and perform two-sample statistical tests for each dimension *d*.

#### 2.6.1 Effect size

For each latent feature *k* and dimension *d* (either clinical or genetic), we compute the effect size Δ_*kd*_ as:

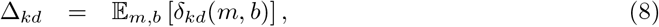

where ***δ***_*kd*_ = {*δ_kd_*(*m, b*)}_∀*m,b*_ is an *M* × *B* matrix of effect sizes for each posterior sample *m* and bootstrap iteration *b*. The expectation is done across all posterior samples and bootstrapping iterations, which are equally probable. For each feature *k* and input dimension *d*, we check for mean differences, i.e.,

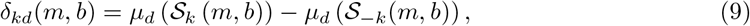

where 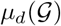 is the mean value of variable *d* for a given subpopulation 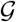.

#### 2.6.2 Statistical significance

To measure how significant an effect size *δ_kd_*(*m, b*) is, for each posterior sample *m* and bootstrap instance *b*, we compute a statistical significance value *ν_kd_*(*m, b*) as the *p*-value resulting from a Fisher test, which is a standard test for discrete variables [50]. We define the *K* × *D* matrix of statistical significance **ϒ**, for each latent feature *k* and input dimension *d* as the median *p*-value across the *M* samples and *B* bootstrapping instances:

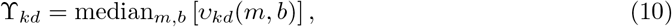

where *ν_kd_* denote the *M* × *B* matrix of statistical significance values *ν_kd_*(*m, b*) for each posterior sample *m* and bootstrapping instance *b*. Finally, we follow the Benjamini Hochberg procedure for multiple hypothesis testing to adjust the statistical significance threshold *α_s_* such that a certain false discovery rate (FDR) is guaranteed [6]. An input dimension *d* (either clinical or genetic) is said to be statistically significant for latent feature *k* if its significance value ϒ_*kd*_ (the median *p*-value across posterior samples and bootstrapping instances) is smaller than the adjusted threshold, i.e., ϒ_*kd*_ < *α_s_*. The whole procedure is summarized in Algorithm 1.

##### Algorithm 1 Statistical approach for discovery of clinico-genetic associations (post-processing procedure).

~~~
**Require:** *M* posterior samples from **Z**, where *P* is the number of subpopulations, and *K* is the number of inferred latent features.
 1: **for** *m* = 1,…,*M* **do**
 2: bootstrap for each subpopulation *B* times
 3: **end for**
 4: **for** *k* = 1,…,*K* **do**
 5: compute effect size according to Eq. 8 and 9.
 6: compute statistical significance (*p*-value) according to the Fisher test adjusting for multiple hypothesis testing [6].
 7: **end for
Ensure:** effect size matrix **Δ** and significance matrix **ϒ**, both of dimensions *K* × *D*
~~~

### 2.7 Experimental Setup

We compare the proposed H-PFA approach with a LMM and a standard case-control set-up for each potential clinico-genetic association. The model parameters for each LMM are found by maximizing the log likelihood using standard optimization techniques within a python platform called LIMIX [30]. In the final step, we obtain *p*-values for each pair (***y***_*q*_, ***x***_*g*_) using likelihood ratio tests. Regarding the case-control analysis, for each clinical term we consider a case and control group corresponding to the patients having that clinical term active or inactive respectively.

Given such a partition, we perform an individual Fisher test for each gene. For all methods, we correct for multiple hypothesis testing based on the Benjamini-Hochberg approach [6]. To quantify the statistical significance of the features discovered by the H-PFA model, we follow the statistical procedure described in Section 2.6. To increase interpretability of the H-PFA model, we force one of the latent features to be active for all patients. This is a common practice in BNP models to capture mean effects [41, 47, 46], which in this case corresponds to phenotypical attributes common to all types of cancer. Finally, we have set the hyperparameters of the proposed H-PFA as *α*_**A**_ = 0.01 and *μ*_A_ = 1, while we infer the values for the concentration parameter *α*.

## 3 Results

### 3.1 Identification of Clinico-Genetic Associations

Figure 2 represents the number of associations found by each method, and how many overlap across techniques. LMM found 14 clinico-genetic associations, CC found 178, and H-PFA found 95.^5^ LMM finds the least number of associations, since it corrects for the cancer type as a confounder effect, and only gets less well-known associations that are present across all types of cancer. CC discovered the highest number of associations, from which only 30% are shared with H-PFA. Out of the 95 associations discovered by the H-PFA approach, 63% were also present in any of the other methods. Figure 3 lists the associations that are shared across methods. Tables 1 and 2 present the list of clinico-genetic associations found by LMM and CC methods (for CC, we only report a random selection of associations, but the complete list can be found in the Appendix).

**Figure 2:**
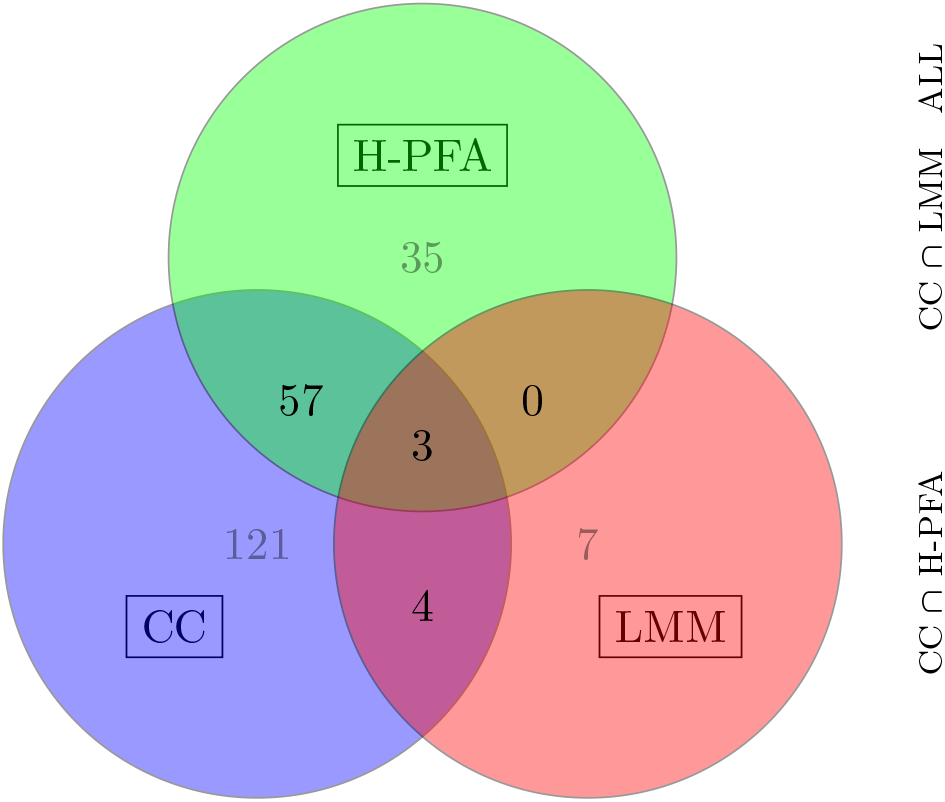
Venn Diagram of number of associations.

**Figure 3:**
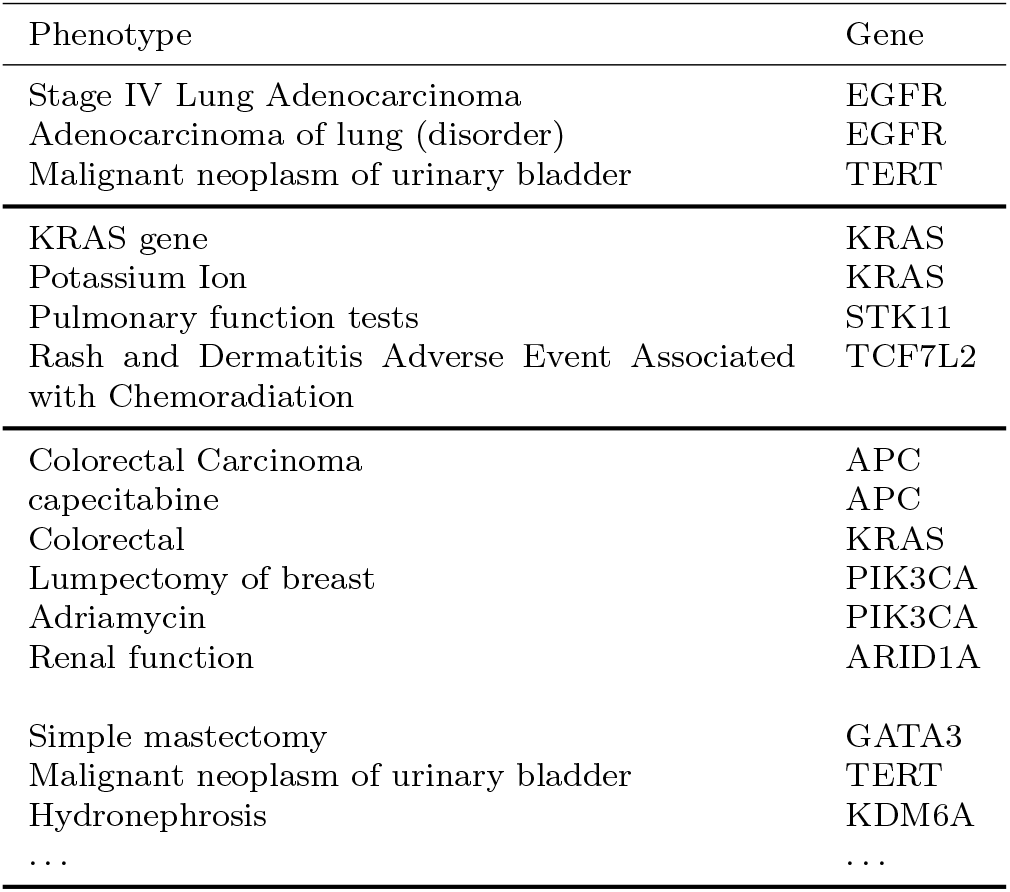
Shared associations across methods.

**Table 1:** Subset of clinico-genetic associations found using the CC setup. A complete list can be found in the Appendix.

| Phenotype | Gene | $\beta_{qg}$ | $p$-value |
| --- | --- | --- | --- |
| pump (device) | APC | 0.61 | 2.78e-53 |
| S3 (sacral segmental innervation) | TP53 | 0.29 | 1.02e-09 |
| Stage IV Lung Adenocarcinoma | EGFR | 0.32 | 2.07e-29 |
| Folinic Acid-Fluorouracil-Irinotecan Regimen | APC | 0.59 | 1.08e-60 |
| Folinic Acid-Fluorouracil-Irinotecan Regimen | KRAS | 0.30 | 1.16e-18 |
| Hepatectomy | APC | 0.65 | 1.21e-52 |
| Hepatectomy | KRAS | 0.27 | 7.27e-12 |
| FOLFOX Regimen | APC | 0.66 | 5.12e-113 |
| Tract | ARID1A | 0.21 | 5.61e-12 |
| Malignant neoplasm of urinary bladder | TERT | 0.55 | 1.07e-61 |
| Renal function | TERT | 0.28 | 8.40e-12 |
| Flushing | APC | 0.30 | 1.54e-11 |
| Non-Small Cell Lung Carcinoma | EGFR | 0.18 | 4.11e-11 |
| Colorectal Carcinoma | APC | 0.61 | 2.34e-63 |
| Adenocarcinoma of lung (disorder) | EGFR | 0.26 | 1.49e-22 |
| Simple mastectomy | PIK3CA | 0.16 | 3.40e-08 |
| Immunotherapy | TERT | 0.23 | 2.76e-12 |
| Imodium | APC | 0.30 | 6.51e-15 |
| capecitabine | APC | 0.30 | 1.94e-15 |
| Pulmonary function tests | STK11 | 0.16 | 2.19e-09 |

**Table 2:** Clinico-genetic associations found using the LMM approach. The associations have been sorted according to the effect size *β_qg_* which refers to the linear weight of the regression, as described in Section 2.4.

| Phenotype | Gene | $\beta_{qg}$ | $p$-value |
| --- | --- | --- | --- |
| Stage IV Lung Adenocarcinoma | EGFR | 0.14 | 9.90e-14 |
| Pulmonary function tests | STK11 | 0.11 | 5.67e-06 |
| Esophagogastroduodenoscopy | ERBB3 | 0.10 | 4.62e-06 |
| Adenocarcinoma of lung (disorder) | EGFR | 0.09 | 3.20e-06 |
| Rash and Dermatitis Adverse Event Associated with Chemoradiation | TCF7L2 | 0.09 | 1.57e-04 |
| Atrophic | PTEN | 0.09 | 2.06e-05 |
| Esophagogastroduodenoscopy | ALK | 0.09 | 1.43e-05 |
| Stage level 2 | ERBB4 | 0.08 | 3.34e-05 |
| Positive Surgical Margin | EP300 | 0.08 | 1.43e-04 |
| Esophagogastroduodenoscopy | CDH1 | 0.08 | 1.33e-04 |
| Esophagogastroduodenoscopy | FLT4 | 0.08 | 4.15e-05 |
| Malignant neoplasm of urinary bladder | TERT | 0.08 | 7.91e-05 |
| Potassium Ion | KRAS | 0.08 | 6.72e-06 |
| KRAS gene | KRAS | 0.07 | 9.76e-05 |

Next, Table 3 shows the list of inferred latent features by the H-PFA model. The bias term F0 reflects the high rate mutation of the TP53 gene which occur across all types of cancer. The TP53 gene is essential for the production of a protein called tumor protein p53. This protein acts as a tumor suppressor, which means that it regulates cell division by keeping cells from growing and dividing too fast or in an uncontrolled way. Because p53 is essential for regulating cell division and preventing tumor formation, it has been nicknamed the “guardian of the genome” [34]. On top of the bias term F0, H-PFA inferred 19 other latent features. Features F3, F5, and F17 capture complex phenotypes (no somatic mutations involved), whereas F4 and F18 mostly capture somatic mutations. Interestingly, F18 relates Esophagogastroduodenoscopy (a test to examine the lining of the esophagus, stomach, and the beginning of the small intestine) to multiple somatic mutations, which was already revealed by LMM in Table 2. The remaining 14 features capture co-ocurrence of somatic mutations and clinical UMLS terms. Some latent features reflect well known relationships in oncology research. To name a few, mutations of gene PIK3CA (captured by F1 and F16) are present in over one-third of breast cancers; such mutations are nowadays known to be oncogenic and also implicated in cervical cancers [18]. Somatic mutations in the triad APC-KRAS-TP53 genes (captured by F0 and F6 together) are prominent in colon cancer [1]. Finally, previous studies have found direct physiological and molecular evidence for a role of gene FOXA1 in controlling cell proliferation in prostate cancer [23], which is accounted for in factor F12.

**Table 3:** Latent features inferred by H-PFA. We depict the UMLS terms and genes with highest weights separately, up until the weight decays more than 50%. *m_k_* is the number of patients with each feature active.

| Feat. | $m_k$ | Phenotypes | Genes |
| --- | --- | --- | --- |
| F0. | 1946 | None | TP53 (0.40) |
| F1. | 460 | Simple mastectomy (0.17), Xeloda (0.15), Lumpectomy of breast (0.12), capecitabine (0.09) | PIK3CA (0.31) |
| F2. | 402 | Renal function (0.29), Coronary Artery Disease (0.28), Stent, device (0.22), cardiologist (0.21), Urology (0.20), Hydronephrosis (0.16) | MTOR (0.04) |
| F3. | 400 | Invasive Ductal Breast Carcinoma (0.57), axillary lymph node dissection (0.38), Simple mastectomy (0.37), Noninfiltrating Intraductal Carcinoma (0.36), Lumpectomy of breast (0.33), Adriamycin (0.29) | None |
| F4. | 392 | None | ETV6 (0.22), PT-PRD (0.19), ATR (0.19), PTPRT (0.17), , ... |
| F5. | 361 | Entire intercostal space (0.22), Midclavicular line (0.21), Per Minute (0.20), Prednisone (0.19), Upper Extremity (0.19), Entire head (0.18), Dizziness (0.17), Redness (0.17), Serum (0.17), Bedtime (qualifier value) (0.16), ... | None |
| F6. | 352 | Colorectal (0.39), FOLFOX Regimen (0.29) | APC (0.71), KRAS (0.47) |
| F7. | 350 | Lobectomy (0.48), Pulmonary function tests (0.29), Thoracotomy (0.27), Non-Small Cell Lung Carcinoma (0.27) | EGFR (0.09) |
| F8. | 326 | Non-Small Cell Lung Carcinoma (0.13), Stage IV Lung Adenocarcinoma (0.10), natural daughter - RoleCode (0.09), pemetrexed (0.08) | KRAS (0.36), STK11 (0.26), KEAP1 (0.20) |
| F9. | 326 | Lytic lesion (0.29), Zometa (0.27), Fracture (0.25), Sclerosis (0.25), Bone Lesion (0.23), Bone structure of sacrum (0.23), Hip arthralgia (0.23), Bone structure of ilium (0.19), Palliative Care (0.17) | ATRX (0.04) |
| F10. | 266 | Stage IV Lung Adenocarcinoma (0.62), pemetrexed (0.61), Adenocarcinoma of lung (disorder) (0.60), mediastinal lymphadenopathy (0.35) | EGFR (0.30), TP53 (0.18) |
| F11. | 265 | FOLFOX Regimen (0.54), KRAS gene (0.45), Folinic Acid-Fluorouracil-Irinotecan Regimen (0.43), Leucovorin (0.43), irinotecan (0.39), Colorectal Carcinoma (0.39), Cold intolerance (0.37), Midclavicular line (0.27), Sigmoid colon (0.27), Colorectal (0.27) | PTPRT (0.05), CARD11 (0.04) |
| F12. | 262 | Prostate carcinoma (0.74), adenocarcinoma of the prostate (0.69), Biopsy of prostate (0.62), Extracapsular (0.55), Lupron (0.46), Personal Attribute (0.40) | FOXA1 (0.11), APC (0.06) |
| F13. | 261 | Tract (0.52), Malignant neoplasm of urinary bladder (0.51), Gross hematuria (0.44), Incontinence (0.29), Immunotherapy (0.28) | TERT (0.66), KDM6A (0.36) |
| F14. | 169 | Lovenox (0.28), Pulmonary Embolism (0.27), Deep Vein Thrombosis (0.23), swollen feet/legs (0.19) | None |
| F15. | 159 | Rectum (0.45), Rash and Dermatitis Adverse Event Associated with Chemoradiation (0.28), capecitabine (0.26), Node stage N0 (0.23) | APC (0.14), TCF7L2 (0.14), TSC2 (0.09) |
| F16. | 149 | Consistency (0.55), Vagina (0.50), Clinic / Center - Mobile (0.43), Bilateral Salpingectomy with Oophorectomy (0.39), Atrophic (0.39), New medications (0.35), Personal Attribute (0.32), Uterus (0.31), Ovarian (0.30), Bone Mineral Density Test (0.30), Ovary (0.29) | PIK3CA (0.12), PTEN (0.07) |
| F17. | 107 | Depression motion (0.95), Structure of long bone (0.82), S3 (sacral segmental innervation) (0.82), pump (device) (0.79), intrahepatic (0.78), Pulse taking (0.74), Midclavicular line (0.73), Entire intercostal space (0.70), Hepatectomy (0.62), Flowcharts (Computer) (0.57) | None |
| F18. | 95 | Esophagogastroduodenoscopy (0.12) | POLE (0.65), ROS1 (0.61), DNMT1 (0.59), ATR (0.58), ATM (0.57), FAT1 (0.54), ... |
| F19. | 41 | Optic Nerve (0.90), Gross hematuria (0.90), Dyspepsia (0.57), Lupron (0.45) | AR (0.22) |

Figure 4 depicts the cancer-specific activation weights 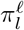 for each type of cancer *ℓ*, as described in previous section.The activation of features present strong variations across cancer types. Some features are clearly cancer-specific (F1 and F3 typically activate for breast carcinoma patients; F6, F11 and F15 are typically active for colorectal cancer; F7, F8 and F10 are almost exclusively active for non-small cell lung cancer, etc.), whereas other factors occur in similar proportions across cancers, e.g., feature F5 which capture typical adverse effects that manifest for all types of cancer (Prednisone is a synthetic corticosteroid drug which is regularly used to treat certain types of cancer, but has significant adverse effects).

**Figure 4:**
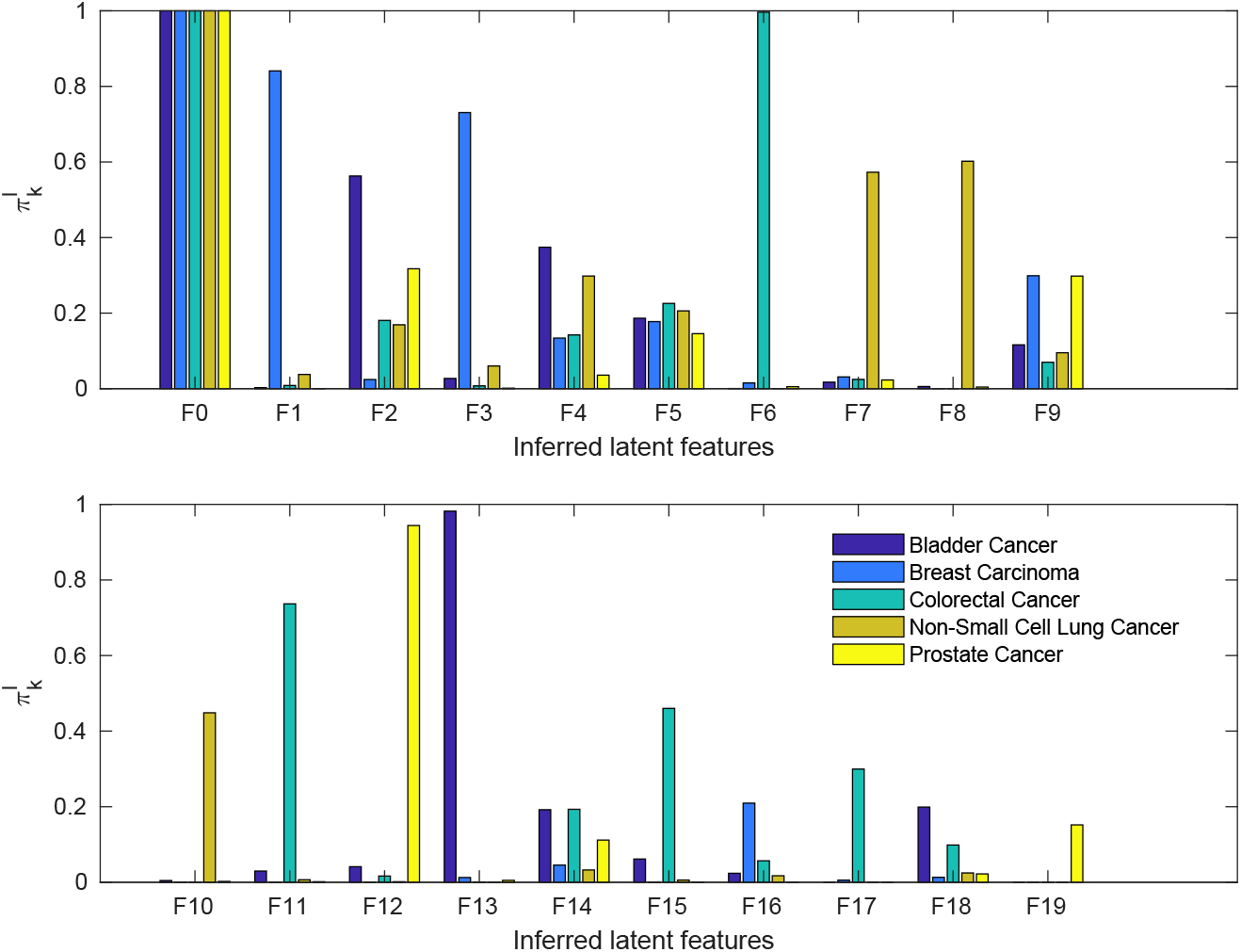
Activation weights 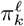 for each cancer type *ℓ* inferred by H-PFA.

Tables 4, 5, 7, 8, and 9 show statistically-significant group associations across different somatic mutations and UMLS terms. For each clinical and genetic term, we give both the effect size and significance. Our method is able to provide concise grouping of both clinical terms and somatic mutations. Among the clinical terms, we find both phenotypical terms, as well as names of chemotherapy medications (Adriamycin, Irinotecan, or Leucovorin). Table 4 shows cancer-specific clinico-genetic associations. We recover well-known associations (such as APC gene mutation being prominant in colorectal cancer, or STK11 to lung carcinoma), but other associations are more surprising, such as GATA3 gene with bone mineral density.

**Table 4:**
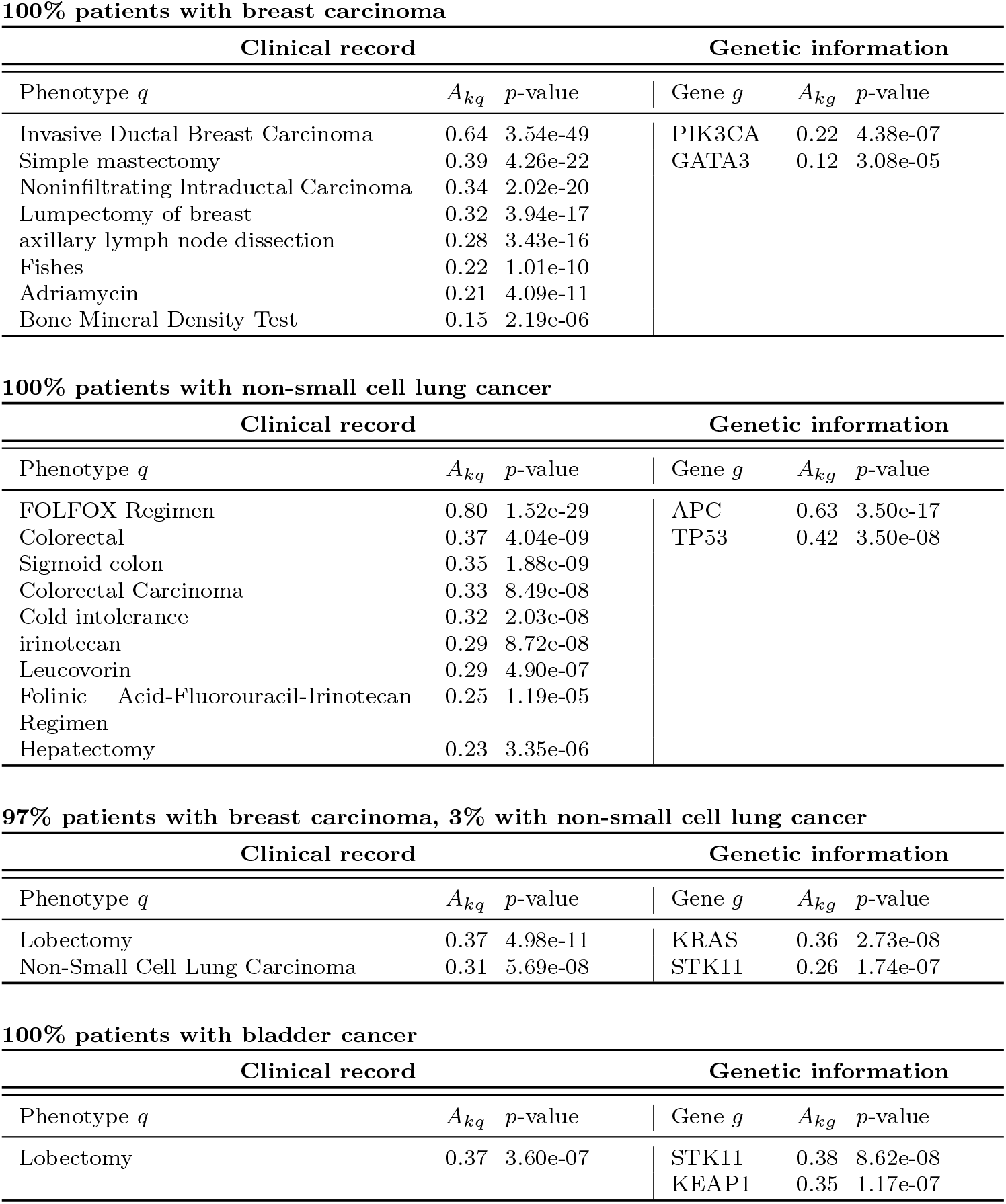
Clinico-genetic associations found by the H-PFA (1/2). These tables depict the statistically significant clinical and genetic features associated to the latent factors listed in Table 3, after applying the statistical methodology described in Section 2.6. On top of each table, we describe the distribution of cancer types of patients for which the association is active.

**Table 5:** Clinico-genetic associations found by the H-PFA (2/2). All these associations involve gene TERT. One same set (depicted in bold) appears in all associations.

| 100% patients with colorectal cancer |  |  |  |  |  |
| --- | --- | --- | --- | --- | --- |
| Clinical record |  |  | Genetic information |  |  |
| Phenotype $q$ | $A_{kq}$ | $p$-value | Gene $g$ | $A_{kg}$ | $p$-value |
| **Tract** | **0.37** | **3.39e-08** | **TERT** | **0.57** | **4.56e-12** |
| **Malignant neoplasm of urinary bladder** | **0.36** | **3.92e-07** | FGFR3 | 0.50 | 4.15e-13 |
| **Gross hematuria** | **0.33** | **2.67e-06** |  |  |  |
| 100% patients with prostate cancer |  |  |  |  |  |
| Clinical record |  |  | Genetic information |  |  |
| Phenotype $q$ | $A_{kq}$ | $p$-value | Gene $g$ | $A_{kg}$ | $p$-value |
| **Gross hematuria** | **0.74** | **1.81e-20** | **TERT** | **0.58** | **5.59e-12** |
| **Malignant neoplasm of urinary bladder** | **0.41** | **2.80e-08** | KDM6A | 0.31 | 6.48e-06 |
| **Tract** | **0.39** | **2.08e-08** |  |  |  |
| chain of objects | 0.32 | 1.92e-06 |  |  |  |
| Hydronephrosis | 0.26 | 2.52e-05 |  |  |  |
| 100% patients with non-small cell lung cancer |  |  |  |  |  |
| Clinical record |  |  | Genetic information |  |  |
| Phenotype $q$ | $A_{kq}$ | $p$-value | Gene $g$ | $A_{kg}$ | $p$-value |
| **Gross hematuria** | **0.39** | **1.72e-07** | **TERT** | **0.64** | **4.86e-13** |
| **Malignant neoplasm of urinary bladder** | **0.37** | **9.85e-07** | FGFR3 | 0.50 | 1.97e-11 |
| **Tract** | **0.30** | **2.19e-05** | KDM6A | 0.36 | 1.20e-06 |
|  |  |  | CREBBP | 0.32 | 3.19e-06 |
| 100% patients with bladder cancer |  |  |  |  |  |
| Clinical record |  |  | Genetic information |  |  |
| Phenotype $q$ | $A_{kq}$ | $p$-value | Gene $g$ | $A_{kg}$ | $p$-value |
| **Malignant neoplasm of urinary bladder** | **0.56** | **4.24e-12** | **TERT** | **0.67** | **1.79e-14** |
| **Tract** | **0.46** | **2.00e-10** | ARID1A | 0.45 | 2.32e-08 |
| Renal function | 0.45 | 5.51e-11 |  |  |  |
| **Gross hematuria** | **0.36** | **8.69e-07** |  |  |  |

Finally, H-PFA found several statistically significant sets of associations involving somatic mutations in gene TERT, as shown in Table 5. Somatic mutations in the gene promoter of telomerase reverse transcriptase (TERT) have been found in 70-79% of bladder tumors in a multi-institutional study published in European Urology [36]. Table 5 shows that TERT mutations are associated to not only malignant neoplasm of urinary bladder (which is not surprising), but also hematuria and hydronephrosis. Hematuria refers to the presence of red blood cells in the urine. Also, hydronephrosis is a condition that typically occurs when the kidney swells due to the failure of normal drainage of urine from the kidney to the bladder. Hydronephrosis is not a primary disease, but results from some other underlying disease (cancer in this case) as the result of a blockage or obstruction in the urinary tract. H-PFA points out to interesting gene relationships (KDM6A, CREBBP, and ARID1A genes) with TERT, which have been partially studied in the literature [33, 39, 25].

## 4 Conclusion

This paper proposes a novel Bayesian nonparametric approach for discovering clinico-genetic associations between somatic mutations and EHR-based clinical features. We present a hierarchical Bernoulli process Poisson factor analysis model based on a hierarchical construction of Beta processes and Bernoulli processes. Our approach is not specific to cancer data nor Electronic Health Records, but can be broadly used to discover associations between arbitrary count data features. Compared to other approaches, our model delivers group-associations instead of pairwise ones, accounting for epistatic and pleiotropical effects straightforwardly. The delivered associations are statistically significant after correction for multiple hypothesis testing combined with a bootstrapping procedure, to better account for false positives. These associations give potentially interesting insights for future research in oncology. Under the proposed model, we hopefully open the door to find new associations that give rise to hypotheses, and if those are validated, then we may get new insights about cancer biology. Ultimately, studies like this one have the potential to lead us towards more accurate diagnosis, and inform us about actionable pathways when considering cancer therapy, where interventions through drug administration can be designed.

## Acknowledgements

This study was supported by the MSK Cancer Center Support Grant (P30 CA008748). M.F.P. was supported by the European Union 7th Framework Programme through the Marie Curie Initial Training Network “Machine Learning for Personalized Medicine” MLPM2012, Grant No. 316861 (to F.P.C and G.R.). This work was also partially supported by MINECO/FEDER (‘ADVENTURE’, id. TEC2015-69868-C2-1-R), and Comunidad de Madrid (project ‘CASI-CAM-CM’, id. S2013/ICE-2845). We gratefully acknowledge helpful discussions with Theofanis Karaletsos, Chris Sander, and Niki Schultz. Moreover, we would like to thank Iker Huerga, Chris Crosbie, Stuart Gardos and David Artz for supporting the work with data deliveries from the clinical data warehouse.

## 5 Appendix: Complete List of Associations

### 5.1 Case-control setup (CC)

**Table 6:**
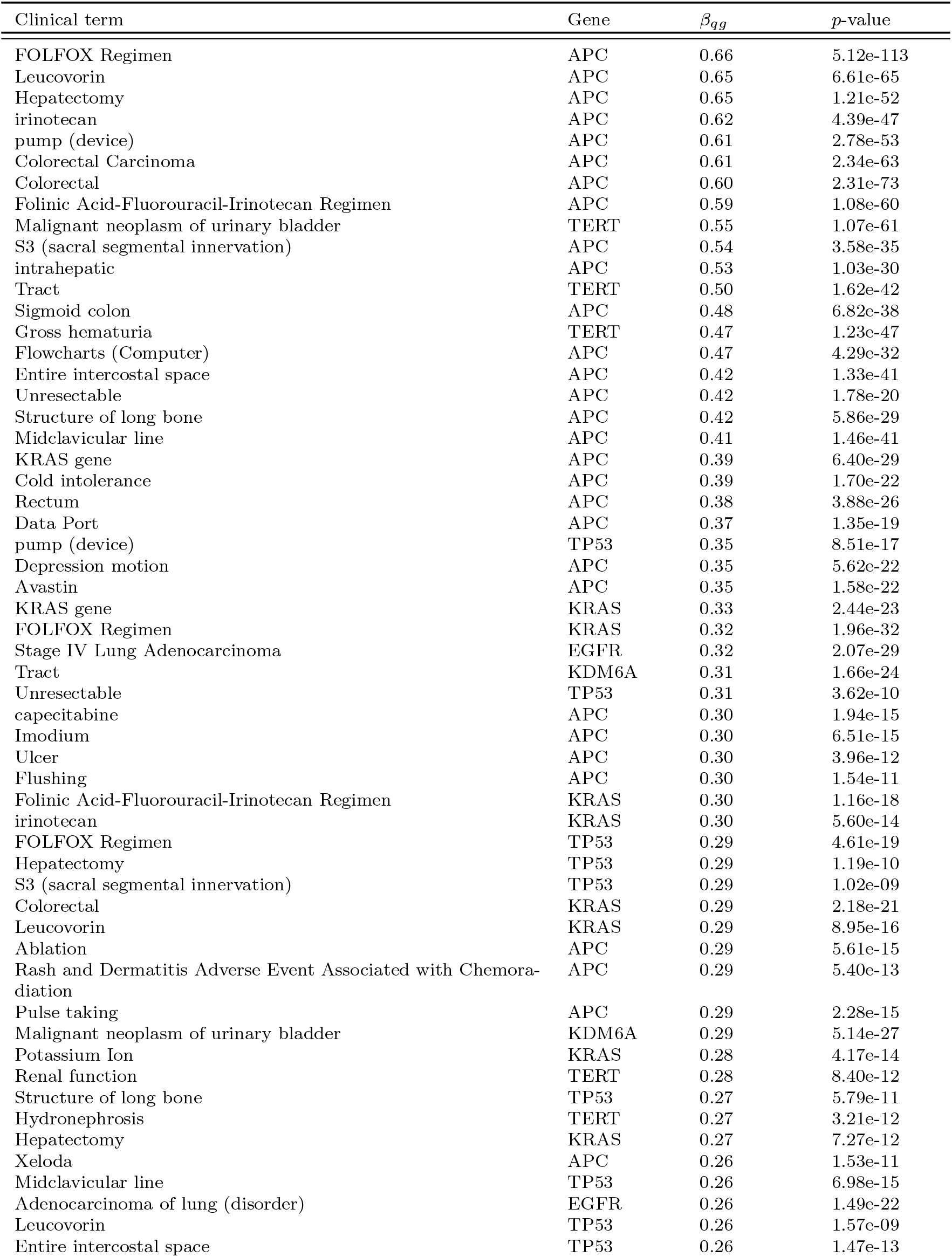
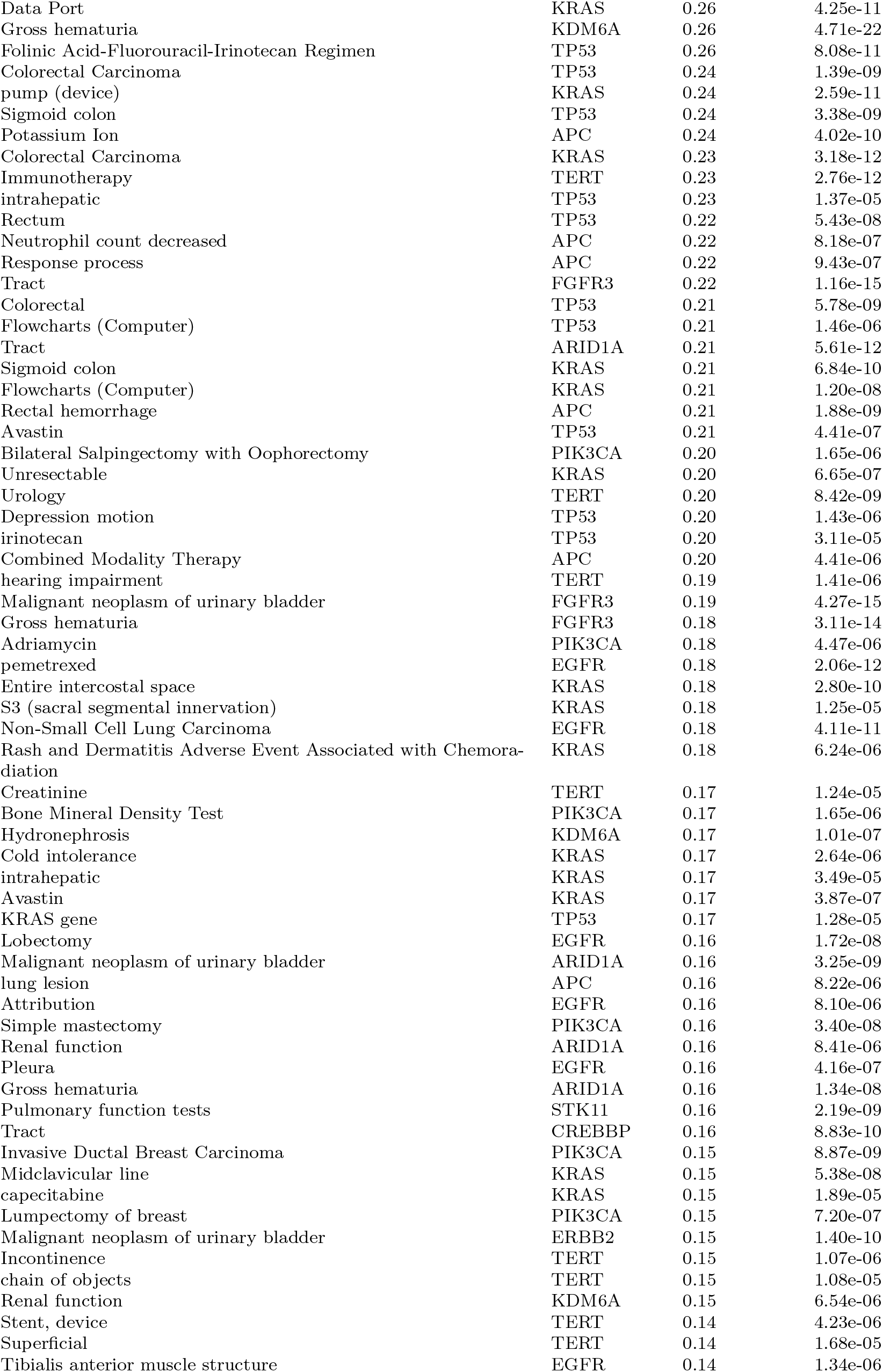
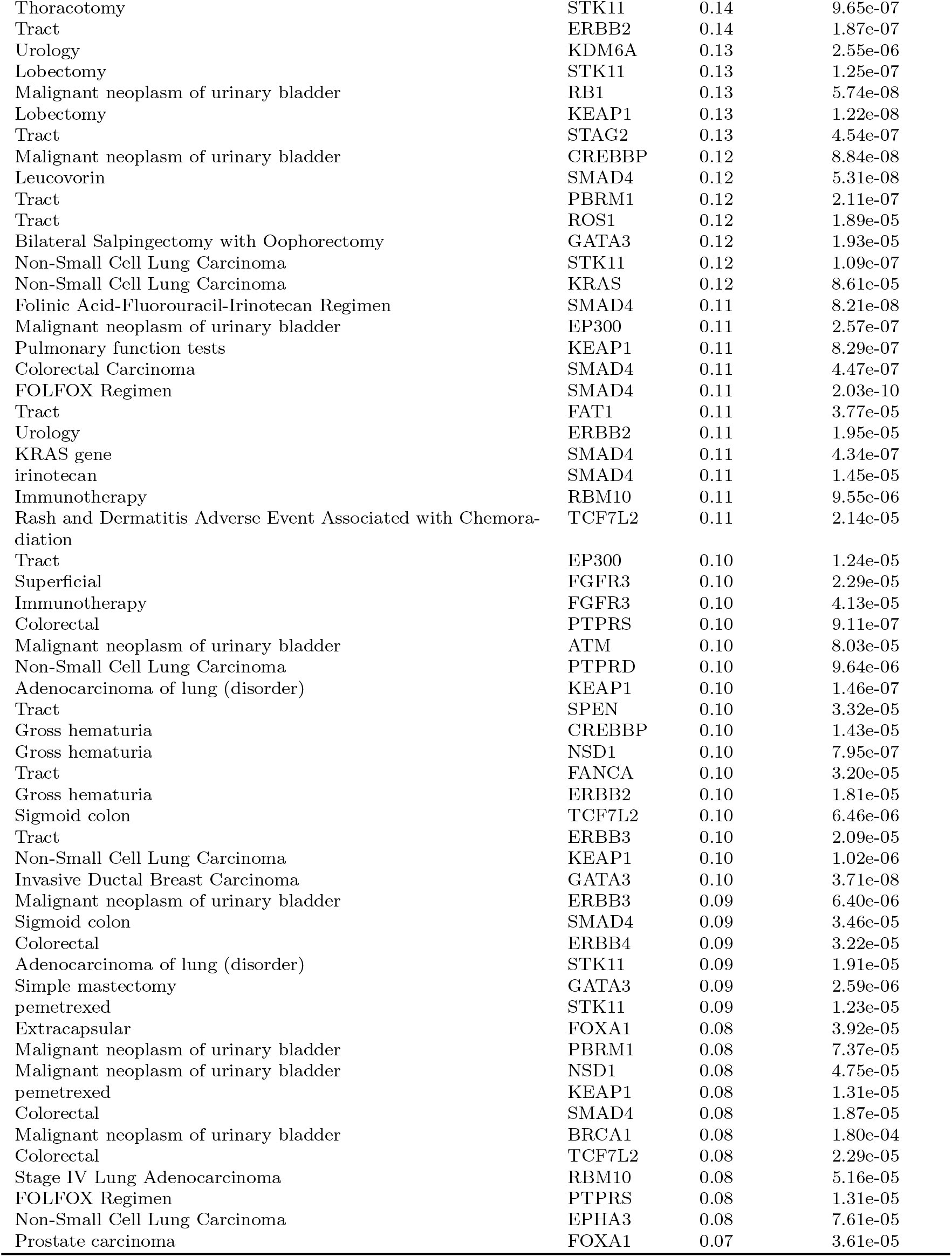
Complete list of clinico-genetic associations found using the Case-Control Set-up. *β_qg_* refers to the linear weight as described in Section 2.4. Associations in bold have also been discovered by the H-PFA.

| Clinical term | Gene | $\beta_{gg}$ | $p$-value |
| --- | --- | --- | --- |
| FOLFOX Regimen | APC | 0.66 | 5.12e-113 |
| Leucovorin | APC | 0.65 | 6.61e-65 |
| Hepatectomy | APC | 0.65 | 1.21e-52 |
| irinotecan | APC | 0.62 | 4.39e-47 |
| pump (device) | APC | 0.61 | 2.78e-53 |
| Colorectal Carcinoma | APC | 0.61 | 2.34e-63 |
| Colorectal | APC | 0.60 | 2.31e-73 |
| Folinic Acid-Fluorouracil-Irinotecan Regimen | APC | 0.59 | 1.08e-60 |
| Malignant neoplasm of urinary bladder | TERT | 0.55 | 1.07e-61 |
| S3 (sacral segmental innervation) | APC | 0.54 | 3.58e-35 |
| intrahepatic | APC | 0.53 | 1.03e-30 |
| Tract | TERT | 0.50 | 1.62e-42 |
| Sigmoid colon | APC | 0.48 | 6.82e-38 |
| Gross hematuria | TERT | 0.47 | 1.23e-47 |
| Flowcharts (Computer) | APC | 0.47 | 4.29e-32 |
| Entire intercostal space | APC | 0.42 | 1.33e-41 |
| Unresectable | APC | 0.42 | 1.78e-20 |
| Structure of long bone | APC | 0.42 | 5.86e-29 |
| Midclavicular line | APC | 0.41 | 1.46e-41 |
| KRAS gene | APC | 0.39 | 6.40e-29 |
| Cold intolerance | APC | 0.39 | 1.70e-22 |
| Rectum | APC | 0.38 | 3.88e-26 |
| Data Port | APC | 0.37 | 1.35e-19 |
| pump (device) | TP53 | 0.35 | 8.51e-17 |
| Depression motion | APC | 0.35 | 5.62e-22 |
| Avastin | APC | 0.35 | 1.58e-22 |
| KRAS gene | KRAS | 0.33 | 2.44e-23 |
| FOLFOX Regimen | KRAS | 0.32 | 1.96e-32 |
| Stage IV Lung Adenocarcinoma | EGFR | 0.32 | 2.07e-29 |
| Tract | KDM6A | 0.31 | 1.66e-24 |
| Unresectable | TP53 | 0.31 | 3.62e-10 |
| capecitabine | APC | 0.30 | 1.94e-15 |
| Imodium | APC | 0.30 | 6.51e-15 |
| Ulcer | APC | 0.30 | 3.96e-12 |
| Flushing | APC | 0.30 | 1.54e-11 |
| Folinic Acid-Fluorouracil-Irinotecan Regimen | KRAS | 0.30 | 1.16e-18 |
| irinotecan | KRAS | 0.30 | 5.60e-14 |
| FOLFOX Regimen | TP53 | 0.29 | 4.61e-19 |
| Hepatectomy | TP53 | 0.29 | 1.19e-10 |
| S3 (sacral segmental innervation) | TP53 | 0.29 | 1.02e-09 |
| Colorectal | KRAS | 0.29 | 2.18e-21 |
| Leucovorin | KRAS | 0.29 | 8.95e-16 |
| Ablation | APC | 0.29 | 5.61e-15 |
| Rash and Dermatitis Adverse Event Associated with Chemora-<br>diation | APC | 0.29 | 5.40e-13 |
| Pulse taking | APC | 0.29 | 2.28e-15 |
| Malignant neoplasm of urinary bladder | KDM6A | 0.29 | 5.14e-27 |
| Potassium Ion | KRAS | 0.28 | 4.17e-14 |
| Renal function | TERT | 0.28 | 8.40e-12 |
| Structure of long bone | TP53 | 0.27 | 5.79e-11 |
| Hydronephrosis | TERT | 0.27 | 3.21e-12 |
| Hepatectomy | KRAS | 0.27 | 7.27e-12 |
| Xeloda | APC | 0.26 | 1.53e-11 |
| Midclavicular line | TP53 | 0.26 | 6.98e-15 |
| Adenocarcinoma of lung (disorder) | EGFR | 0.26 | 1.49e-22 |
| Leucovorin | TP53 | 0.26 | 1.57e-09 |
| Entire intercostal space | TP53 | 0.26 | 1.47e-13 |
| Data Port | KRAS | 0.26 | 4.25e-11 |
| Gross hematuria | KDM6A | 0.26 | 4.71e-22 |
| Folinic Acid-Fluorouracil-Irinotecan Regimen | TP53 | 0.26 | 8.08e-11 |
| Colorectal Carcinoma | TP53 | 0.24 | 1.39e-09 |
| pump (device) | KRAS | 0.24 | 2.59e-11 |
| Sigmoid colon | TP53 | 0.24 | 3.38e-09 |
| Potassium Ion | APC | 0.24 | 4.02e-10 |
| Colorectal Carcinoma | KRAS | 0.23 | 3.18e-12 |
| Immunotherapy | TERT | 0.23 | 2.76e-12 |
| intrahepatic | TP53 | 0.23 | 1.37e-05 |
| Rectum | TP53 | 0.22 | 5.43e-08 |
| Neutrophil count decreased | APC | 0.22 | 8.18e-07 |
| Response process | APC | 0.22 | 9.43e-07 |
| Tract | FGFR3 | 0.22 | 1.16e-15 |
| Colorectal | TP53 | 0.21 | 5.78e-09 |
| Flowcharts (Computer) | TP53 | 0.21 | 1.46e-06 |
| Tract | ARID1A | 0.21 | 5.61e-12 |
| Sigmoid colon | KRAS | 0.21 | 6.84e-10 |
| Flowcharts (Computer) | KRAS | 0.21 | 1.20e-08 |
| Rectal hemorrhage | APC | 0.21 | 1.88e-09 |
| Avastin | TP53 | 0.21 | 4.41e-07 |
| Bilateral Salpingectomy with Oophorectomy | PIK3CA | 0.20 | 1.65e-06 |
| Unresectable | KRAS | 0.20 | 6.65e-07 |
| Urology | TERT | 0.20 | 8.42e-09 |
| Depression motion | TP53 | 0.20 | 1.43e-06 |
| irinotecan | TP53 | 0.20 | 3.11e-05 |
| Combined Modality Therapy | APC | 0.20 | 4.41e-06 |
| hearing impairment | TERT | 0.19 | 1.41e-06 |
| Malignant neoplasm of urinary bladder | FGFR3 | 0.19 | 4.27e-15 |
| Gross hematuria | FGFR3 | 0.18 | 3.11e-14 |
| Adriamycin | PIK3CA | 0.18 | 4.47e-06 |
| pemetrexed | EGFR | 0.18 | 2.06e-12 |
| Entire intercostal space | KRAS | 0.18 | 2.80e-10 |
| S3 (sacral segmental innervation) | KRAS | 0.18 | 1.25e-05 |
| Non-Small Cell Lung Carcinoma | EGFR | 0.18 | 4.11e-11 |
| Rash and Dermatitis Adverse Event Associated with Chemora-<br>diation | KRAS | 0.18 | 6.24e-06 |
| Creatinine | TERT | 0.17 | 1.24e-05 |
| Bone Mineral Density Test | PIK3CA | 0.17 | 1.65e-06 |
| Hydronephrosis | KDM6A | 0.17 | 1.01e-07 |
| Cold intolerance | KRAS | 0.17 | 2.64e-06 |
| intrahepatic | KRAS | 0.17 | 3.49e-05 |
| Avastin | KRAS | 0.17 | 3.87e-07 |
| KRAS gene | TP53 | 0.17 | 1.28e-05 |
| Lobectomy | EGFR | 0.16 | 1.72e-08 |
| Malignant neoplasm of urinary bladder | ARID1A | 0.16 | 3.25e-09 |
| lung lesion | APC | 0.16 | 8.22e-06 |
| Attribution | EGFR | 0.16 | 8.10e-06 |
| Simple mastectomy | PIK3CA | 0.16 | 3.40e-08 |
| Renal function | ARID1A | 0.16 | 8.41e-06 |
| Pleura | EGFR | 0.16 | 4.16e-07 |
| Gross hematuria | ARID1A | 0.16 | 1.34e-08 |
| Pulmonary function tests | STK11 | 0.16 | 2.19e-09 |
| Tract | CREBBP | 0.16 | 8.83e-10 |
| Invasive Ductal Breast Carcinoma | PIK3CA | 0.15 | 8.87e-09 |
| Midclavicular line | KRAS | 0.15 | 5.38e-08 |
| capecitabine | KRAS | 0.15 | 1.89e-05 |
| Lumpectomy of breast | PIK3CA | 0.15 | 7.20e-07 |
| Malignant neoplasm of urinary bladder | ERBB2 | 0.15 | 1.40e-10 |
| Incontinence | TERT | 0.15 | 1.07e-06 |
| chain of objects | TERT | 0.15 | 1.08e-05 |
| Renal function | KDM6A | 0.15 | 6.54e-06 |
| Stent, device | TERT | 0.14 | 4.23e-06 |
| Superficial | TERT | 0.14 | 1.68e-05 |
| Tibialis anterior muscle structure | EGFR | 0.14 | 1.34e-06 |
| Thoracotomy | STK11 | 0.14 | 9.65e-07 |
| Tract | ERBB2 | 0.14 | 1.87e-07 |
| Urology | KDM6A | 0.13 | 2.55e-06 |
| Lobectomy | STK11 | 0.13 | 1.25e-07 |
| Malignant neoplasm of urinary bladder | RB1 | 0.13 | 5.74e-08 |
| Lobectomy | KEAP1 | 0.13 | 1.22e-08 |
| Tract | STAG2 | 0.13 | 4.54e-07 |
| Malignant neoplasm of urinary bladder | CREBBP | 0.12 | 8.84e-08 |
| Leucovorin | SMAD4 | 0.12 | 5.31e-08 |
| Tract | PBRM1 | 0.12 | 2.11e-07 |
| Tract | ROS1 | 0.12 | 1.89e-05 |
| Bilateral Salpingectomy with Oophorectomy | GATA3 | 0.12 | 1.93e-05 |
| Non-Small Cell Lung Carcinoma | STK11 | 0.12 | 1.09e-07 |
| Non-Small Cell Lung Carcinoma | KRAS | 0.12 | 8.61e-05 |
| Folinic Acid-Fluorouracil-Irinotecan Regimen | SMAD4 | 0.11 | 8.21e-08 |
| Malignant neoplasm of urinary bladder | EP300 | 0.11 | 2.57e-07 |
| Pulmonary function tests | KEAP1 | 0.11 | 8.29e-07 |
| Colorectal Carcinoma | SMAD4 | 0.11 | 4.47e-07 |
| FOLFOX Regimen | SMAD4 | 0.11 | 2.03e-10 |
| Tract | FAT1 | 0.11 | 3.77e-05 |
| Urology | ERBB2 | 0.11 | 1.95e-05 |
| KRAS gene | SMAD4 | 0.11 | 4.34e-07 |
| irinotecan | SMAD4 | 0.11 | 1.45e-05 |
| Immunotherapy | RBM10 | 0.11 | 9.55e-06 |
| Rash and Dermatitis Adverse Event Associated with Chemora-<br>diation | TCF7L2 | 0.11 | 2.14e-05 |
| Tract | EP300 | 0.10 | 1.24e-05 |
| Superficial | FGFR3 | 0.10 | 2.29e-05 |
| Immunotherapy | FGFR3 | 0.10 | 4.13e-05 |
| Colorectal | PTPRS | 0.10 | 9.11e-07 |
| Malignant neoplasm of urinary bladder | ATM | 0.10 | 8.03e-05 |
| Non-Small Cell Lung Carcinoma | PTPRD | 0.10 | 9.64e-06 |
| Adenocarcinoma of lung (disorder) | KEAP1 | 0.10 | 1.46e-07 |
| Tract | SPEN | 0.10 | 3.32e-05 |
| Gross hematuria | CREBBP | 0.10 | 1.43e-05 |
| Gross hematuria | NSD1 | 0.10 | 7.95e-07 |
| Tract | FANCA | 0.10 | 3.20e-05 |
| Gross hematuria | ERBB2 | 0.10 | 1.81e-05 |
| Sigmoid colon | TCF7L2 | 0.10 | 6.46e-06 |
| Tract | ERBB3 | 0.10 | 2.09e-05 |
| Non-Small Cell Lung Carcinoma | KEAP1 | 0.10 | 1.02e-06 |
| Invasive Ductal Breast Carcinoma | GATA3 | 0.10 | 3.71e-08 |
| Malignant neoplasm of urinary bladder | ERBB3 | 0.09 | 6.40e-06 |
| Sigmoid colon | SMAD4 | 0.09 | 3.46e-05 |
| Colorectal | ERBB4 | 0.09 | 3.22e-05 |
| Adenocarcinoma of lung (disorder) | STK11 | 0.09 | 1.91e-05 |
| Simple mastectomy | GATA3 | 0.09 | 2.59e-06 |
| pemetrexed | STK11 | 0.09 | 1.23e-05 |
| Extracapsular | FOXA1 | 0.08 | 3.92e-05 |
| Malignant neoplasm of urinary bladder | PBRM1 | 0.08 | 7.37e-05 |
| Malignant neoplasm of urinary bladder | NSD1 | 0.08 | 4.75e-05 |
| pemetrexed | KEAP1 | 0.08 | 1.31e-05 |
| Colorectal | SMAD4 | 0.08 | 1.87e-05 |
| Malignant neoplasm of urinary bladder | BRCA1 | 0.08 | 1.80e-04 |
| Colorectal | TCF7L2 | 0.08 | 2.29e-05 |
| Stage IV Lung Adenocarcinoma | RBM10 | 0.08 | 5.16e-05 |
| FOLFOX Regimen | PTPRS | 0.08 | 1.31e-05 |
| Non-Small Cell Lung Carcinoma | EPHA3 | 0.08 | 7.61e-05 |
| Prostate carcinoma | FOXA1 | 0.07 | 3.61e-05 |

### 5.2 Hierarchical Poisson Factor Analysis

**Table 7:**
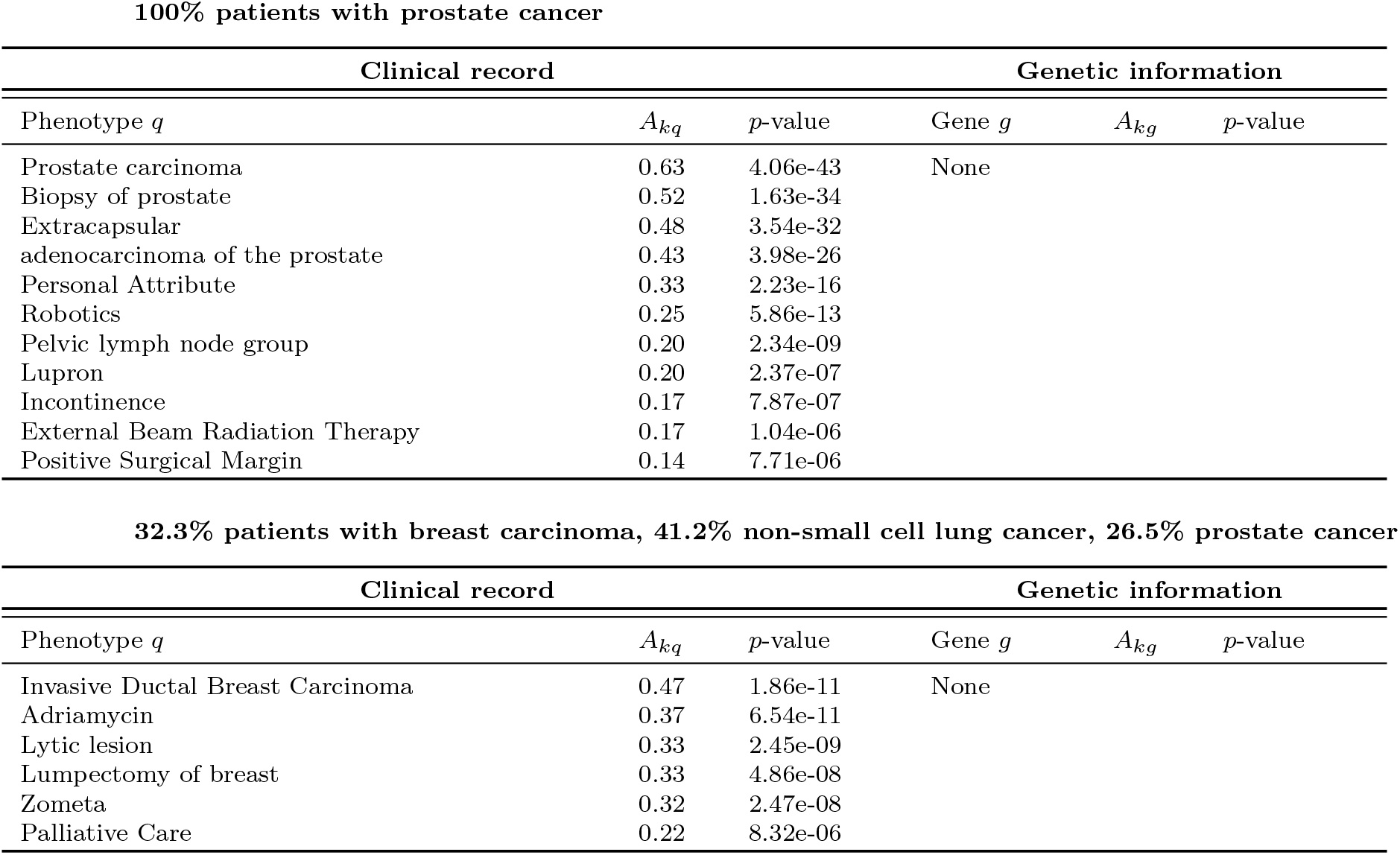
Additional clinical associations (complex phenotypes) found by the H-PFA.

**Table 8:** Additional clinico-genetic association found by the H-PFA involving APC gene. This group of associations was found in a subgroup of 100% bladder cancer patients.

| Clinical record |  |  | Genetic information |  |  |
| --- | --- | --- | --- | --- | --- |
| Phenotype $q$ | $A_{kq}$ | $p$-value | Gene $g$ | $A_{kg}$ | $p$-value |
| FOLFOX Regimen | 0.77 | 3.47e-19 | APC | 0.59 | 5.34e-11 |
| Rectum | 0.59 | 1.56e-14 |  |  |  |
| Sigmoid colon | 0.43 | 8.36e-09 |  |  |  |
| Rash and Dermatitis Adverse Event Associated with Chemoradiation | 0.42 | 1.24e-09 |  |  |  |
| capecitabine | 0.42 | 5.37e-09 |  |  |  |
| Colorectal | 0.42 | 1.19e-07 |  |  |  |
| Folinic Acid-Fluorouracil-Irinotecan Regimen | 0.40 | 1.11e-07 |  |  |  |
| KRAS gene | 0.38 | 2.12e-07 |  |  |  |
| Ulcer | 0.36 | 3.64e-08 |  |  |  |
| irinotecan | 0.35 | 3.10e-07 |  |  |  |
| Rectal hemorrhage | 0.34 | 3.31e-06 |  |  |  |
| Avastin | 0.32 | 1.13e-05 |  |  |  |
| Leucovorin | 0.29 | 2.95e-05 |  |  |  |
| Combined Modality Therapy | 0.26 | 5.02e-05 |  |  |  |
| Response to treatment | 0.25 | 4.10e-05 |  |  |  |

**Table 9:** Additional clinico-genetic associations found by the H-PFA involving EGFR gene. This group of associations was found in a subgroup of 100% non-small cell lung cancer patients.

| Clinical record |  |  | Genetic information |  |  |
| --- | --- | --- | --- | --- | --- |
| Phenotype $q$ | $A_{kq}$ | $p$-value | Gene $g$ | $A_{kg}$ | $p$-value |
| Stage IV Lung Adenocarcinoma | 0.64 | 3.47e-15 | EGFR | 0.35 | 8.65e-06 |
| Adenocarcinoma of lung (disorder) | 0.56 | 3.92e-11 |  |  |  |
| pemetrexed | 0.51 | 7.89e-10 |  |  |  |
| Tibialis anterior muscle structure | 0.32 | 3.40e-06 |  |  |  |

## 6 Appendix: Inference details for Poisson Factor Analysis

Poisson factorization models have been successfully applied for recommendation systems [15], topic modeling [14], and analysis of Electronic Health Records among others [21]. Let 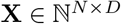 be a sparse matrix of count-data observations with *N* samples and *D* dimensions. The generative model for the Bernoulli process Poisson factor analysis (PFA) is given by:

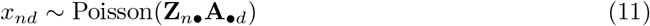

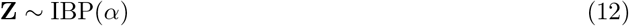

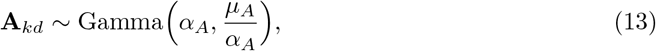

where **Z** is a *N* × *K* matrix of binary weights, and **A** is a *K* × *D* matrix of non-negative hidden factors. Direct inference in such models is intractable, but we can easily solve the problem using MCMC techniques. For each observation x_nd_, we introduce the auxiliary variables 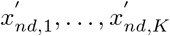 such that 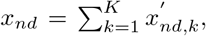, and 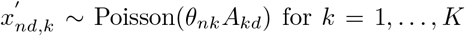. Each Poisson count is separated in a sum of Poisson contributions corresponding to each latent factor. Given such auxiliary variables, the model is conditionally conjugate, and a Gibbs sampler can be derived straightforwardly. In particular, we use the following theorem:

### Theorem 1

*Let Y*_1_,…,*Y_n_ be Poisson distributed random variables with rates* λ_1_,…,λ_*n*_ *respectively. Let us define* 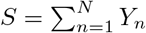. *Then*,

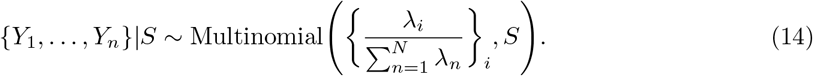

Using Theorem 1, 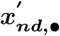 can be sampled from a Multinomial given *x_nd_*, ***θ_n•_*** and **A_•*d*_**. In the following, we propose two MCMC algorithms: a collapsed Gibbs sampler where matrix **A** is marginalized out using a Laplace approximation, and an uncollapsed slice sampler version which allows for parallel sampling of both the elements in **Z** and **A** given the auxiliary variables 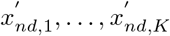. The results shown in this work have been generated using the uncollapsed Gibbs Sampler.

### 6.1 Collapsed Gibbs Sampler

We first propose a collapsed Gibbs sampler where matrix **A** is marginalized out, and we only need to sample the elements of matrix **Z.** We need to compute its posterior distribution:

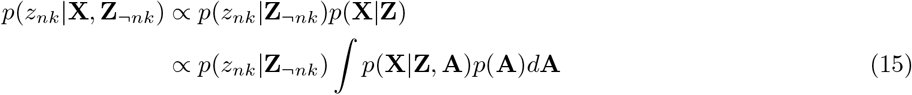

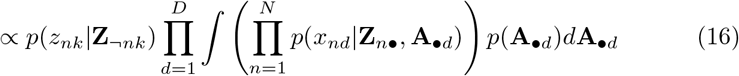

In order to approximate the integral in (16), we resort to a Laplace approximation, which assumes that:

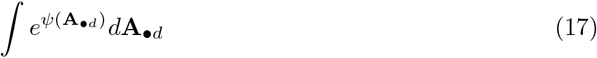

has a peak at a certain value of 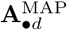. The idea is to Taylor-expand the un-normalized log-posterior of **A_•*d*_** and approximate *e*^*ψ*(**A_•*d*_**)^ by an unnormalized Gaussian. The integral thus corresponds to the normalizing constant of this Gaussian, in our case:

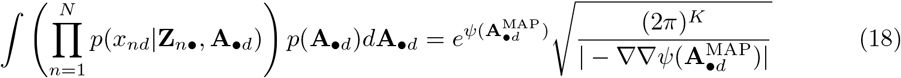

**Equations to find maximum a posteriori** 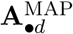. Let us define *ψ*(**A_•*d*_**) as the unnormalized log-posterior of **A_•*d*_**, i.e,

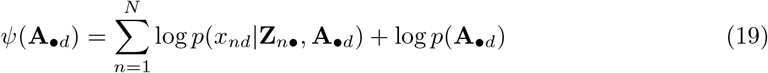

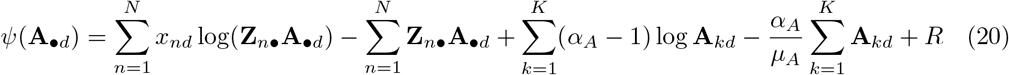

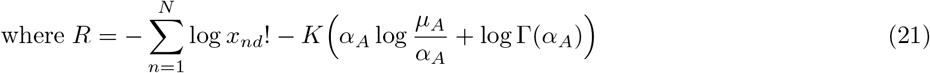

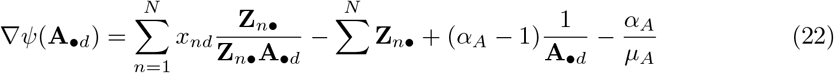

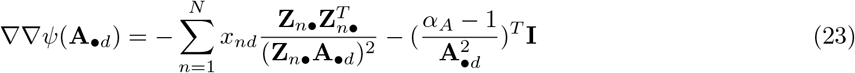

In order to find the maximum value 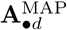, we can use either Newton’s method or gradient descent. Where applicable, Newton’s method might converge faster towards a local maximum or minimum than gradient descent. Newton’s method is an iterative method for optimization where each value 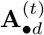 at iteration *t* is computed as:

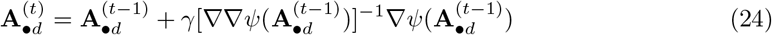

where *γ* ∈ (0,1] is the step-size of the algorithm. Note that for the Laplace approximation to work properly, −∇∇*ψ*(**A_•*d*_**) should be a positive semi-definite matrix. This is guaranteed only if *α_B_* > 1, so the collapsed Gibbs sampler will only work for shape parameters bigger than one, resulting in non-sparse **A** matrices.

### 6.2 Uncollapsed Gibbs Sampler

Inference for the PFA model can be performed using an uncollapsed Gibbs sampler together with a slice sampler for semi-ordered stick-breaking representation of the IBP [44]. For the sake of completeness, the slice sampling procedure for matrix **Z** is described in Algorithm 2. Using the auxiliary random variables described at the beginning of this Appendix, the complete conditionals can be easily derived as follows:

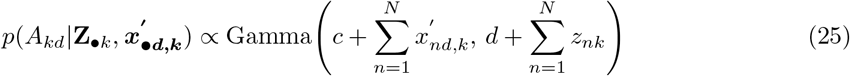

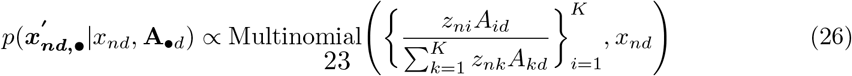

**Algorithm 2.**
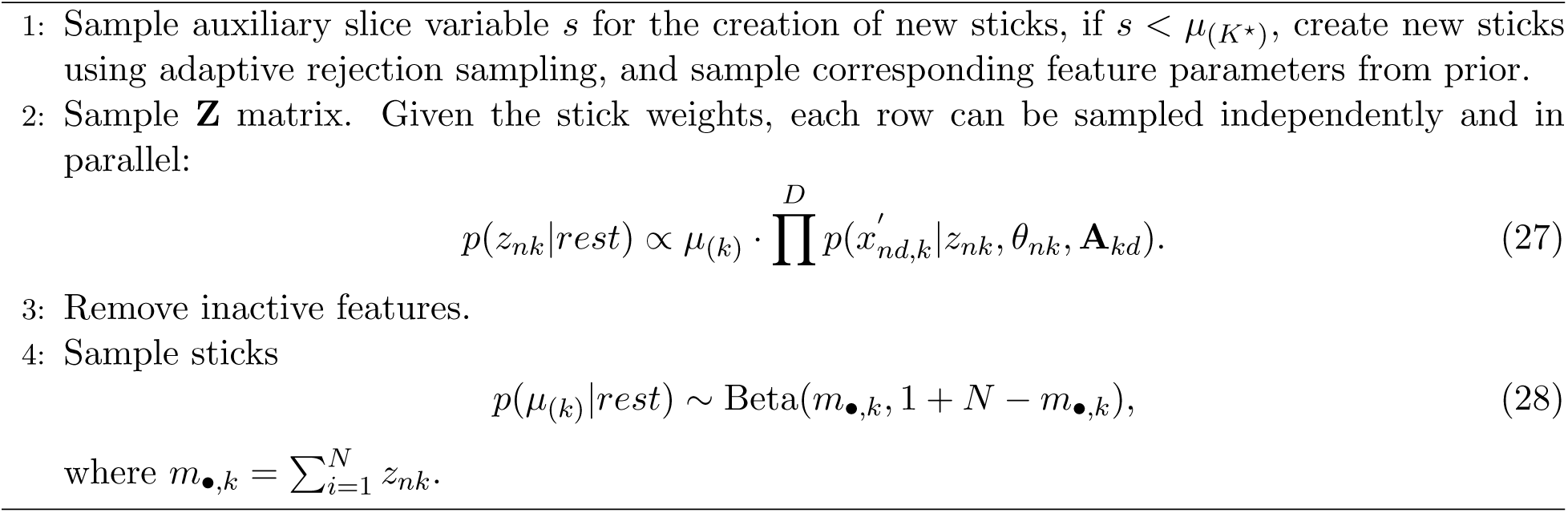
Slice sampler for the semi-ordered stick-breaking representation of the IBP [44].

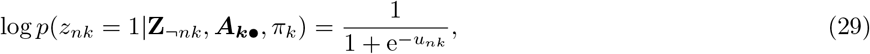

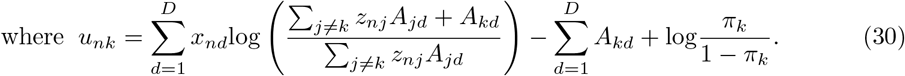

Inference for the hierarchical Poisson Factor Analysis (H-PFA) is analoguous to Algorithm 2 except step 4, where we sample the per-category feature probability 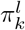 for each cancer type *l* and feature *k* from a Beta distribution based on counts per cancer type as follows:

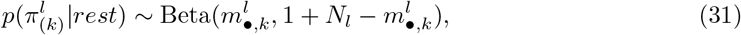

where *N_l_* is the number of patients with cancer type *l*, 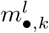 is the number of patients having feature *k* active and cancer type *l, r_n_* is the cancer type indicator for patient *n*, and 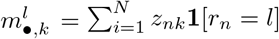.

## Footnotes

1 https://www.mskcc.org/msk-impact

2 The considered sequencing technology is able to remove most of the technical noise, in contrast to other technologies.

3 Source code available at: https://metamap.nlm.nih.gov/

4 Note that statistical significance could also be accounted for using Bayesian factors or posterior predictive checks [13]. We here adopt the most established approach in the field for statistical significance.

5 The H-PFA model is very flexible, as it can also find correlations between the genes, or between the clinical terms. 95 is the number of clinico-genetic associations only.

